# *La révolution de l’ADN*: biocompatible and biosafe DNA data storage

**DOI:** 10.1101/2022.08.25.505104

**Authors:** Alexandre Maes, Jeanne Le Peillet, Achille Julienne, Clémence Blachon, Nicolas Cornille, Mariette Gibier, Erfane Arwani, Zhou Xu, Pierre Crozet, Stéphane D. Lemaire

## Abstract

DNA data storage is an emerging technology that has the potential to replace bulky, fragile and energy-intensive current digital data storage media. Here, we report a storage strategy called DNA Drive, that organizes data on long double stranded replicative DNA molecules. The DNA Drive has unlimited storage capacity, and its encoding scheme ensures the biosafety of the process by limiting the potential of the DNA sequence to code for mRNA and proteins. Using our approach, we encoded two historical texts from the French Revolution, the Declaration of the Rights of Man and of the Citizen of 1789 and the Declaration of the Rights of Woman and of the Female Citizen published in 1791. In contrast to previous DNA storage strategies, the biocompatibility of the DNA Drive enables biological manipulation of the data including low cost copy.

**One-Sentence Summary:** The DNA Drive is a biosafe and biocompatible DNA data storage strategy with unlimited storage capacity.

## Main Text

The storage of digital data is a major challenge in our modern societies. Current digital media stored in data centers are fragile, bulky and energy-intensive. The lifespan of current digital media storage systems (optical media, magnetic tapes, hard disk drives or flash memory) does not exceed seven years on average. Hence, these data must be regularly copied and kept at constant temperature and humidity resulting in a colossal energy cost. Data centers consume 2% of the world’s electricity, and their carbon footprint exceeds that of civil aviation (*1*). The ongoing digital transformation of our societies requires a major disruption of digital storage technologies. DNA is the media used by living organisms to store genetic information. DNA data storage is an emerging technology that is stable for thousands of years, extremely compact and energy-efficient. DNA stability outcompetes any man-made media, as exemplified by the recovery of a near-complete genome from a 1-million-year-old mammoth (*2*). With a maximal density of 4.5 x 10^20^ bytes, *i.e.* 0.45 Zettabytes (ZB) equivalent to 450 x 10^6^ Terabytes (TB) per gram of DNA, all of humanity's digital data in 2019 (45 ZB) could fit in 100 g of DNA (*3*). Moreover, DNA storage does not require energy since these molecules are perfectly stable at room temperature under appropriate conditions (*4, 5*).

The first significant demonstration of DNA data storage was published in 2012 (*6*). Since then a number of studies improved encoding algorithms and barcoding, enabling error correction, direct access and compression (*7–13*). Until now, DNA storage has been carried out mainly using chemically synthesized oligonucleotide pools (short single stranded DNA sequences < 200 bases), which are stored and read *in vitro.* While this approach validated the feasibility of DNA data storage, it has several limitations including a low density and rather limited and expensive editing and copying capabilities. In order to overcome these limitations, we aim at using the biological processes and properties of living organisms to meet the challenges of storing data on DNA.

Our original DNA storage strategy, named DNA Drive, uses long replicative double stranded DNA molecules as a physical support compatible with both *in vitro* and *in vivo* manipulation (Fig.1). Once built, the molecules can be introduced in living organisms, enabling their manipulation using *in vivo* processes, *e.g.* for copy or edition. The DNA Drive allows an ordered physical multi-scale organization of the data on DNA molecules supporting theoretically unlimited drive size, enabling random access and facilitating assembly after sequencing (Fig. 2).

**Fig. 1.**
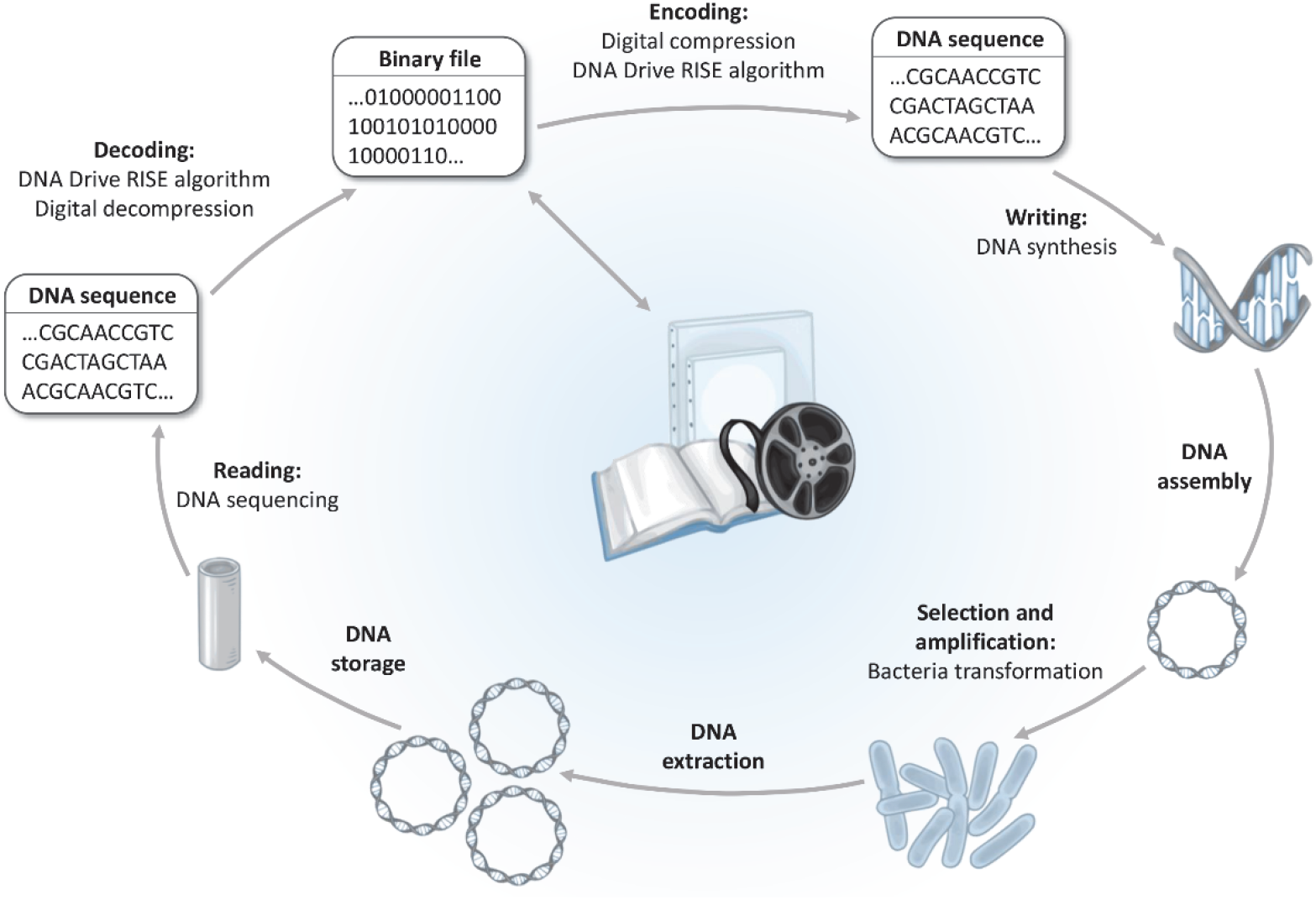
Overall DNA Drive storage process from file to file. The binary file is converted to a biosafe and biocompatible DNA sequence, synthesized, assembled on plasmids, amplified biologically, extracted and stored. Reading of the file is performed by DNA sequencing and conversion of the obtained sequence to the binary file.

**Fig. 2.**
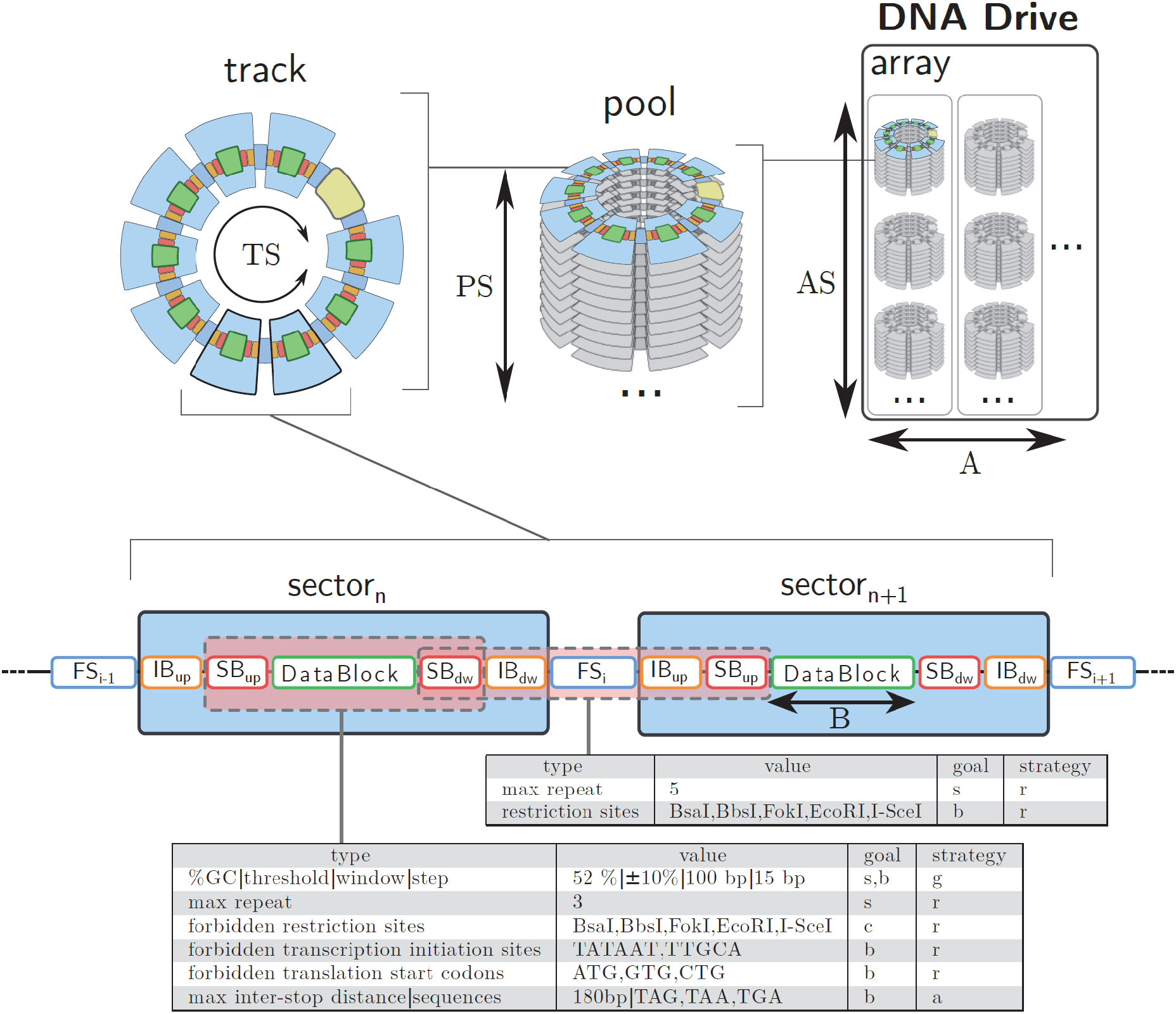
Structure and constraints of the DNA Drive. Binary data are converted to DNA by CGK conversion and stored into DNA Drive datablocks of size B. Each datablock is flanked by two system blocks (SB), two index blocks (IB) and two fusion sites (FS) located upstream (up) or downstream (dw) of the datablock. TS (Track Size) sectors are assembled into replicative molecules called tracks. PS (Pool Size) tracks corresponding to a unique set of index blocks are grouped into a pool. AS (Array Size) pools are physically assembled into A arrays to constitute the DNA Drive. Compatibilization is performed by removing motifs (r), adding motifs (a) or adjusting GC content (g) to ensure optimal DNA synthesis/sequencing (s), DNA construction (c) and biosafety (b).

Several studies reported the storage of data on replicative DNA molecules (*14–17*), but none have considered the possibility that the DNA sequence could encode unwanted biological information as recently demonstrated (*18*). This could represent a risk for the host organism, resulting in data loss, or even a risk for the manipulator or the environment. Accordingly, we developed for the DNA Drive an encoding algorithm that limits the coding potential of the sequence by removal of transcription and translation initiation sequences, and addition of translation termination signals (Fig. 1 and 3). Hence, encoding and assembly are performed using a biocompatible and biosafe algorithm named RISE (Random In-Silico Evolution), which allows conversion of binary data into nucleotide sequences while ensuring the compatibility of the sequence with (i) DNA synthesis and sequencing technologies, (ii) the physical assembly of DNA molecules from isolated data blocks, and (iii) the biosafe manipulation of these biocompatible molecules in living organisms.

**Fig. 3.**
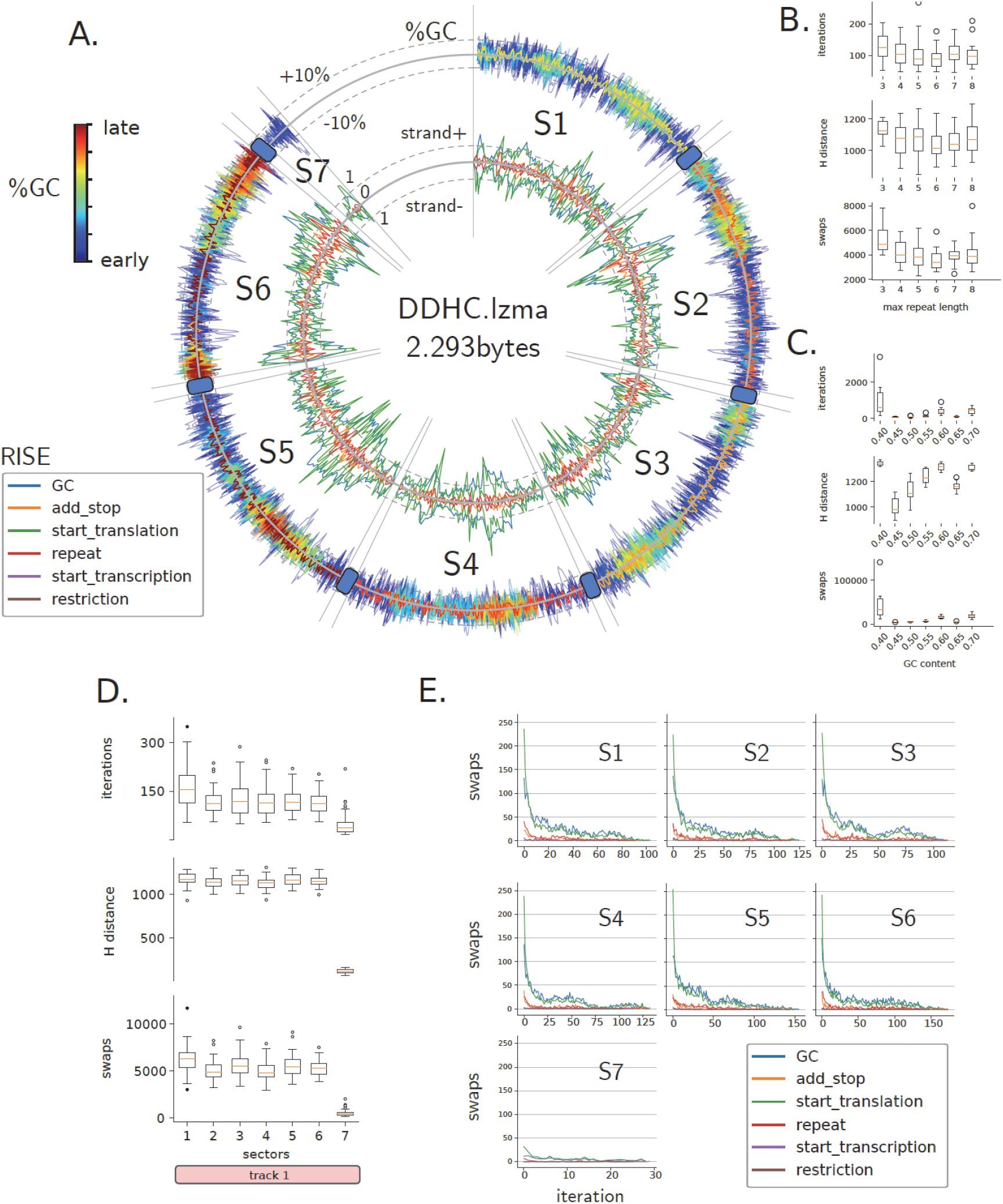
Biosafe and biocompatible encoding of DDHC. Full biocompatibilization of the 7 sectors of DDHC by adding or removing motifs on both strands and adjusting GC content. During RISE, nucleotides can be swapped into their binary synonym without altering digital information. **A.** Number and nature of swap positions (inner circle, mean number of swaps in a sliding window of 30 bp; see legend insert for colors) and GC content evolution (outer circle) during RISE. The color scale indicates whether the swaps for GC content were performed early or late during RISE. **B.** Parameters of RISE simulations of DDHC for various repeat length thresholds (N=40). The Hamming distance (H distance) was computed between the initial and the final fully compatible sequence. Swaps are the number of nucleotide exchanges during all iterations. The cycles indicate the number of RISE iterations. **C.** Parameters of RISE simulations for various final GC content +/− 10 % (N=20). **D.** RISE report for all sectors of track 1 for independent initial CGK random draws (N=40). Note that sector 7 is smaller than the others. **E.** Number of swaps performed for each constraint during iterative evolution of RISE on DDHC. For A, D and E the constraints are described in Fig. 2. The box and whisker plots show the 10th (lower whisker), 25th (base of box), 75th (top of box) and 90th (top whisker) percentiles. Outliers are plotted as individual data points. Hamming (H) distance is computed between the initial and final fully compatible sequence.

Structurally, the DNA Drive is composed of storage units called sectors (Fig. 2). Each sector is composed of a data-block (DB) of adjustable length (B) were the digital information is stored. This DB is flanked by two system blocks (SB, length S), two index blocks (IB, length I), and two fusion sites (FS, length F). Sectors are assembled into circular replicative molecules called tracks, using the flanking fusion sites. The size of a track (TS) is the number of sectors per track. Multiple tracks are pooled together and the pool size (PS) is determined by the number of available index blocks in the DNA Drive. The DNA Drive is sequenced at the pool level to recover data. Pools are physically organized into arrays of size AS, numbered from 1 to A, and their total number is theoretically unlimited. The structure of the DNA Drive is described by the B-I-F-TS-PS-AS-A parameters (Fig. 2).

Each bit of the binary file is converted into DNA using the one bit per base Church-Gao-Kosuri (CGK) encoding scheme (*4*) attributing randomly A or C for 0 and T or G for 1. Combinatorial effect of the CGK method confers flexibility in the conversion of binary information into DNA, allowing to apply multiple constraints on the final sequence, which are necessary for (i) DNA synthesis and sequencing (limitation of homopolymer length, controlled GC content), (ii) DNA assembly (forbidden restriction sites) and (iii) DNA biosafety (limiting transcription and translation). The amount of conversion alternatives (2^n^ DNA sequences for one binary sequence of length n) and the variety of constraints requires a random exploratory method, described in this study as the Random Iterative in-Silico Evolution (RISE).

To demonstrate the efficiency of our DNA storage system, we physically constructed two DNA Drives containing historical founding texts from the French Revolution: “La Déclaration des droits de l’homme et du citoyen” (1789) (The Declaration of the Rights of Man and of the Citizen) and “La Déclaration des droits de la femme et de la citoyenne” (Olympe de Gouges, 1791), *i.e.* the Declaration of the Rights of Woman and of the Female Citizen. The two DNA Drives were assembled and stored dried in metallic minicapsules (DNAshell® from Imagene) ensuring their stability for thousands of years (*4*). The data could be read many times with 100% fidelity.

“La Déclaration des droits de l’homme et du citoyen” (DDHC.txt, 5,253 bytes) and “La Déclaration des droits de la femme et de la citoyenne” (DDFC.txt, 28,763 bytes) were used to illustrate file storage into a DNA Drive of 3.24 Gb, structurally described as B3000S1I25TS9PS10000AS96A1 using the B-I-F-TS-PS-AS-A parameters (Fig. 2). Both files were compressed using the Lempel-Ziv-Markov chain algorithm (LZMA) to limit file size and bit repetitions. Chunks of 375 bytes (3,000 bits) were contiguously allocated to DNA Drive sectors (7 sectors on one track for DDHC.lzma, 2,293 bytes, and 31 sectors on 4 tracks for DDFC.lzma, 11,377 bytes, Fig. S1 and S2 respectively) and converted to DNA by an initial random CGK conversion. However, the DNA sequences generated from the binary file by random CGK draws do not meet the constraints in terms of coding potential, GC content, homopolymers or absence of specific restriction sites (Fig. S3 to S9). The RISE algorithm was designed to iteratively modify the nucleotide sequence and allow convergence towards a biocompatible and biosafe sequence without modification of the binary data (Fig. S10). The RISE constraint parameters employed are detailed in Fig. 2. The coding potential was decreased by removing *E. coli* σ^70^ and σ^54^ −10 motifs to limit RNA production (*19*), by removing translation start codons (ATG, GTG, CTG), and by addition of translation stop codons to yield a DB sequence with at least one stop codon every 180 nucleotides in all 6 reading frames. To limit read and write errors, the GC content was locally adjusted to that of *E. coli* and the maximum homopolymer length was limited to 3 nucleotides. Specific restriction site sequences were also removed to facilitate *in vitro* DNA manipulation.

Practically, the RISE algorithm scans for the presence or absence of specific features, which are then removed or added respectively without altering the binary data, by replacing a nucleotide by its binary synonym, *i.e.* either A by C, or G by T, or C by A, or T by G (Fig. S10). For feature removal, one random nucleotide is swapped in the sequence. For feature addition, a random synonymous sequence is swapped around a seeding position. Because random synonymous modifications can create new incompatible features, multiple iterations are necessary to produce a complete biocompatible sequence that follows the specifications of the DNA Drive (Fig. S11 and S12).

During RISE, hotspots of compatibilization changes are observed along sectors on both strands, with a dominance in GC content adjustment and translation start codon removal (Fig. 3A, Fig. S13). These two constraints are highly predominant during the initial steps of sequence evolution (Fig. 3E). Relaxation of homopolymer length threshold results in a slight increase of RISE efficiency (Fig. 3B). Extreme GC constraints induce a deeper combinatory exploration, resulting in a large increase of RISE cycles and swaps (Fig. S12B). Therefore, GC adjustment is allowed in a large GC window from 40% to 70% GC (Fig. 3C). Multiple RISE simulations on the same initial draw results in various final compatible sequences indicating that random driven processes of RISE allow various paths of compatibilization (Fig. S11). Still, overall compatibilization is homogeneously spread along all the sectors. Successful RISE is reached for all sectors and both files independently of the initial random draw with a mean of 1.84 and 1.78 swaps/base for DDHC and DDFC respectively (Fig. 3, Fig. S11, S12A, S13).

To provide random access to each DB through PCR amplification and facilitate sequence assembly after sequencing, each DB is flanked by a pair of 25-mers barcodes named index blocks (IB, Fig. 2). An ordered collection of 90,000 pairs of IB (Table S1), devoid of homopolymers and forbidden restriction sites, was established by filtering of a previously published library of 240,000 barcodes (*20*). This ordered collection enables random access to any sector in a pool of tracks containing a total of 90,000 DB, for instance a pool size of 10,000 plasmids of 9 sectors. System blocks (SB) are attributed to a random nucleotide, used as a wild card to overcome unsolvable combinations at the junction between data-blocks and ordered barcodes (Fig. 2). Ten 4-mer sequences were used as fusion sites to assemble 9 sectors in a single plasmid. These specifications confer a storage density of 0.98 bit per base.

Each sector was synthesized chemically, assembled and cloned by Twist Bioscience (San Francisco, USA), together with its flanking fusion sites and a BsaI restriction site allowing release of the sector with protruding fusion sites. Each track was assembled using Golden Gate cloning (*21*) in a replicative plasmid, which was amplified in *E. coli* and extracted using DNA maxipreparations to obtain billions of copies of each track. These low-cost copies enabled generation of 250 copies of each DNA Drive, composed of one track for DDHC (50 μg of plasmid DNA, *i.e.* 2500 billion copies of the file per DNA Drive) and 4 tracks for DDFC (5 μg, *i.e.* 100 billion copies per DNA Drive). The DNA of each DNA Drive was lyophilized and encapsulated in DNAShell® metallic minicapsules by Imagene SA (Evry, France) to ensure their conservation at ambient temperatures for thousands of years (*4, 5*). After conservation of the capsules at ambient temperatures for several months (up to 6 months), a capsule was opened, DNA was resuspended in pure water and a fraction was sequenced using an Oxford Nanopore MinION™ sequencer (Oxford Nanopore Technologies, Oxford, UK) (Fig. S14). Basecalled reads were mapped to tracks according to their barcodes. Assembled and polished sequences were then used to generate high accuracy DB sequences. The nucleotide sequence of each DB was converted to binary sequences using the CGK decoding scheme (A=C=0, G=T=1) and the binary file obtained by combining contiguous chunks was then uncompressed with the LZMA algorithm. The original file could be recovered at 100% with no error for each DNA Drive sequenced, demonstrating the robustness of the DNA preservation in DNAShell minicapsules and the efficiency and reliability of the DNA Drive storage strategy. To explore the storage ability of the DNA Drive, experimental and simulated sequencing depth revealed that the B3000S1I25TS9PS10000AS96A1 configuration is suitable for MinION sequencer for complete data recovery (Fig. S14). Future qualitative and quantitative advances in long-read sequencing (basecalling, read length, sequence coverage, assembly and polishing) will allow to continuously increase the storage ability of the DNA Drive pools.

The two capsules containing the DNA Drives of the Declaration of the Rights of Man and of the Citizen (1789) and the Declaration of the Rights of Woman and of the Female Citizen (1791) were registered officially at the French National Archives on November the 23^rd^, 2021. They were deposited in the 1790 safe, “*l’Armoire de fer”*, next to the original paper version of the Declaration of the Rights of Man and of the Citizen from 1789. These two texts are now preserved for thousands of years with no energy input.

## Supporting information

Supplementary files DataS1-S8

## Acknowledgments

The authors would like to thank Bruno Ricard, Françoise Banat-Berger and their colleagues from Archives Nationales for their strong and enthusiastic support for this work. We also thank Dr. Geneviève Fraisse, Pr. Raphaël Ersham, Olivier Blanc, and Marcel Gauchet for their help for digitization of the texts. We thank Marthe Colotte and Sophie Tuffet from Imagene for encapsulation of the DNA Drive. We thank Emily Leproust and Twist Bioscience for synthesis of the DNA Drive fragments. We also thank Claire de Thoisy-Méchin and Marion Valzy for their help with communication. Overall, we thank all the persons that have provided their enthusiastic support for this project.

## Author contributions

Conceptualization: AM, JLP, ZX, SDL, PC

Methodology: AM, JLP, AJ, SDL, PC

Investigation: AM, JLP, AJ, CB, NC, MG, PC

Visualization: AM, JLP, AJ, CB

Funding acquisition: SDL, PC, EA

Project administration: SDL, PC, EA

Supervision: SDL, PC

Writing – original draft: AM, JLP, AJ, SDL, PC

## Data and materials availability

All important data are available in the main text or the supplementary materials. Plasmids and raw data will be made available upon request. Code is available at https://github.com/alexmaes/RISE.

