## Supplementary text, FigS1-S14 for "*La révolution de l’ADN*: biocompatible and biosafe DNA data storage"

**Supplementary Materials**

**Supplementary Text**

**Figs. S1 to S14**

Fig. S1. Map of the DDHC plasmid.
Fig. S2. Maps of the DDFC plasmids
Fig. S3. Distribution of GC content before and after RISE biocompatibilization
Fig. S4. Distribution of homopolymer number and length before RISE biocompatibilization
Fig. S5. Distribution of specific restriction sites before removal by RISE
Fig. S6. Transcription initiation site content before RISE biocompatibilization
Fig. S7. Translation initiation sites before RISE biocompatibilization
Fig. S8. Distance between stop codons before RISE biocompatibilization
Fig. S9. Number and frequency of open reading frames before RISE biocompatibilization
Fig. S10. RISE processing for biocompatibilization
Fig. S11. Distribution of swaps during RISE biocompatibilization of DDHC
Fig. S12. RISE robustness and efficiency
Fig. S13. Biosafe and biocompatible encoding of DDFC
Fig. S14. Nanopore sequencing of the DDHC and DDFC DNA drives

**Table S1**

Table S1. Ordered list of the 90 000 pairs of barcodes required for a 3.28 GB DNA Drive

**Data S1 to S8**

Data S1. DDHC.txt. Text version of DDHC.

Data S2. DDHC.lzma. Compressed version of DDHC.txt.

Data S3. DDHC_sectors.fasta. Biocompatible sectors of DDHC.

Data S4. DDHC_plasmid.fasta. Final sequence of the plasmid containing DDHC.

Data S5. DDHC.txt. Text version of DDFC.

Data S6. DDFC.lzma. Compressed version of DDHC.txt.

Data S7. DDFC_sectors.fasta. Biocompatible sectors of DDFC

Data S8. DDFC_plasmids.fasta. Final sequence of the four plasmids containing DDFC.

**Supplementary Text**

### Materials and Methods

All chemicals were obtained from Sigma-Aldrich, unless otherwise specified.

#### *Encoding*

The two text files containing "la Déclaration des droits de l'homme et du citoyen" (DDHC.txt) and "la Déclaration des droits de la femme et de la citoyenne" (DDFC.txt) were encoded in ISO8859-1 (latin-1) and subsequently compressed with the lossless data compression Lempel-Ziv-Markov chain algorithm (LZMA) using pylzma v0.5.

#### *Conversion and compatibilization*

Conversion from bits to DNA was performed using the Church-Gao-Kosuri (CGK) one bit per base convention, A=C=0, T=G=1 (*6*). The initial random nucleotide draw for each bit was centered on the final expected GC content, [GC]f. For each cycle of Random Iterative in-Silico Evolution (RISE), all bits were sequentially swapped to their CGK counterpart from start to end, to fulfil the constraints and maintain the binary information. For GC content ([GC]) adjustment, in a sliding window (w), if [GC]w>[GC]f*1+threshold, a random G or C in w was swapped to T or A respectively. Conversely, if [GC]w<[GC]f*1-threshold, a random A or T in w was swapped to G or C respectively. The threshold for [GC] variation was fixed at 10 %, based on *in vivo* variations classically observed in *E. coli*. To disrupt a motif, a random position in the motif was swapped to its CGK counterpart. To add a specific motif on each frame of both strands, if dist(motif[i],motif[i+1])>threshold, an available synonym motif was screened and swapped around a random normally distributed in-frame position with μ=dist(motif[i],motif[i+1])/2 and σ=90. The threshold for stop codons frequency was set at 180 bp to avoid any synthesis of peptides > 60 amino acids. New iterations were performed until the complete compatibilization of the sector.

*Barcode library and pre-compatibilization*

A total of 90,000 pairs of barcodes were selected from 240,000 unique orthogonal 25-mers DNA barcode probes (*20*). GC content were already setup at +/-51%. Selection and ordering of barcodes were performed by avoiding specific restriction sites and homopolymers into non-data-block regions (Table S1).

*Transformation, culture and screening of Escherichia coli*

Bacterial growth was performed at 37°C in LB broth supplemented with agar (2% m/V), ampicillin (100 mg/L), kanamycin (50 mg/L) and X-gal (40 mg/L) when required. Chemically competent *E. coli* DH10b (New England Biolabs) were used for transformation (by heat shock following the manufacturer's instructions) and maintenance of plasmids.

To identify positive clones, colony PCR were performed using the Quick-Load R *Taq* 2x reagent (New England Biolabs) according to the manufacturer's recommendations with the corresponding primers.

*Parts/Fragment cloning*

DNA fragments (7 for DDHC, 31 for DDFC,) were synthesized by Twist Bioscience before cloning into a level 0 plasmid (pAGM9121, (*22*)) by Golden Gate Cloning. The tracks were assembled into MoClo plasmids (*23, 24*). The DDHC track (Fig. S1) was assembled in a single Golden Gate cloning step into pAGM9121. To assemble the 3 first tracks of DDFC we first constructed intermediate plasmids containing 3 to 6 sections (Fig. S2A) before assembling the final plasmid from these intermediate fragments (Fig. S2B). The fourth track of DDFC was assembled in a single cloning step as in the case of DDHC. All primers were synthetized by Eurofins Genomics. Molecular biology kits (gel extraction and miniprep kits) were purchased from Macherey-Nagel and used following the manufacturer's instructions. All PCR reactions, unless otherwise stated, were performed using Phusion DNA Polymerase (NEB).

*Golden Gate Assembly*

Restriction-ligation reactions were performed using BbsI-HF (NEB, 20 units per reaction) for level 0 plasmids or Bsal-HFv2 enzyme (NEB, 10 units per reaction) for level 1 (Tracks) cloning, and T4 DNA ligase (NEB, 2000 units per reaction) in CutSmart Buffer (NEB) supplemented with 1 mM ATP (from a 10 mM solution prepared in 100 mM Tris-HCl pH 7.9) in a total volume of 20 µL. All parts were mixed equimolarly in the reaction (20 fmol for level 0 plasmids, 15 fmol for level 1 plasmids). The mixture was incubated for 6 cycles (10 min at 37°C, 10 min at 16°C), followed by 1 hour at 37°C and a last step at 80°C for 20 min. Then 10 units of BsaI-HFv2 were added to the reaction and incubated 12 h at 37°C.

*Quality control and sequencing*

All plasmids were controlled by differential restriction. Level 1 and M plasmids (Tracks) were sequenced using Nanopore sequencing as follows. About 150 fmol of DNA were linearized by EcoRI (NEB) and purified on AMPure XP beads (Beckman-Coulter). DNA libraries were constructed using SQK-LSK109 and NBD104 sequencing kits and protocols (Oxford Nanopore Technologies). Runs of 48h were performed on a MinION and 9.4.1 flow cells. High accuracy base-calling was done using Guppy v.3.4.5. Tracks were then demultiplexed and ordered according to their barcodes before assembling and polishing using either 4 cycles of minimap v2.17-r941/racon v1.4.22 or Flye v2.9, and medaka v1.5 respectively. Using minimap2/racon, the original draft sequence is randomly picked up among the longest reads.

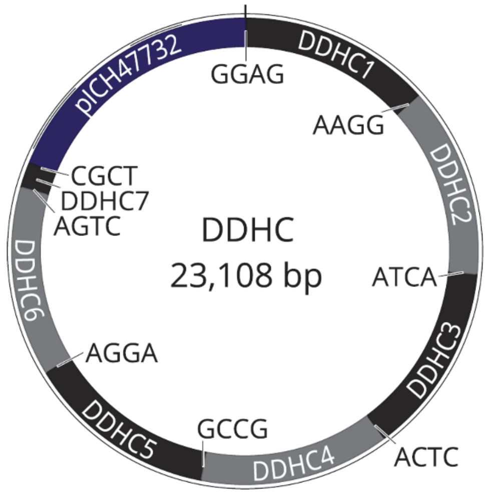


**Fig. S1. Map of the DDHC plasmid.**
Map of the plasmid containing the 7 sectors of the DDHC file.

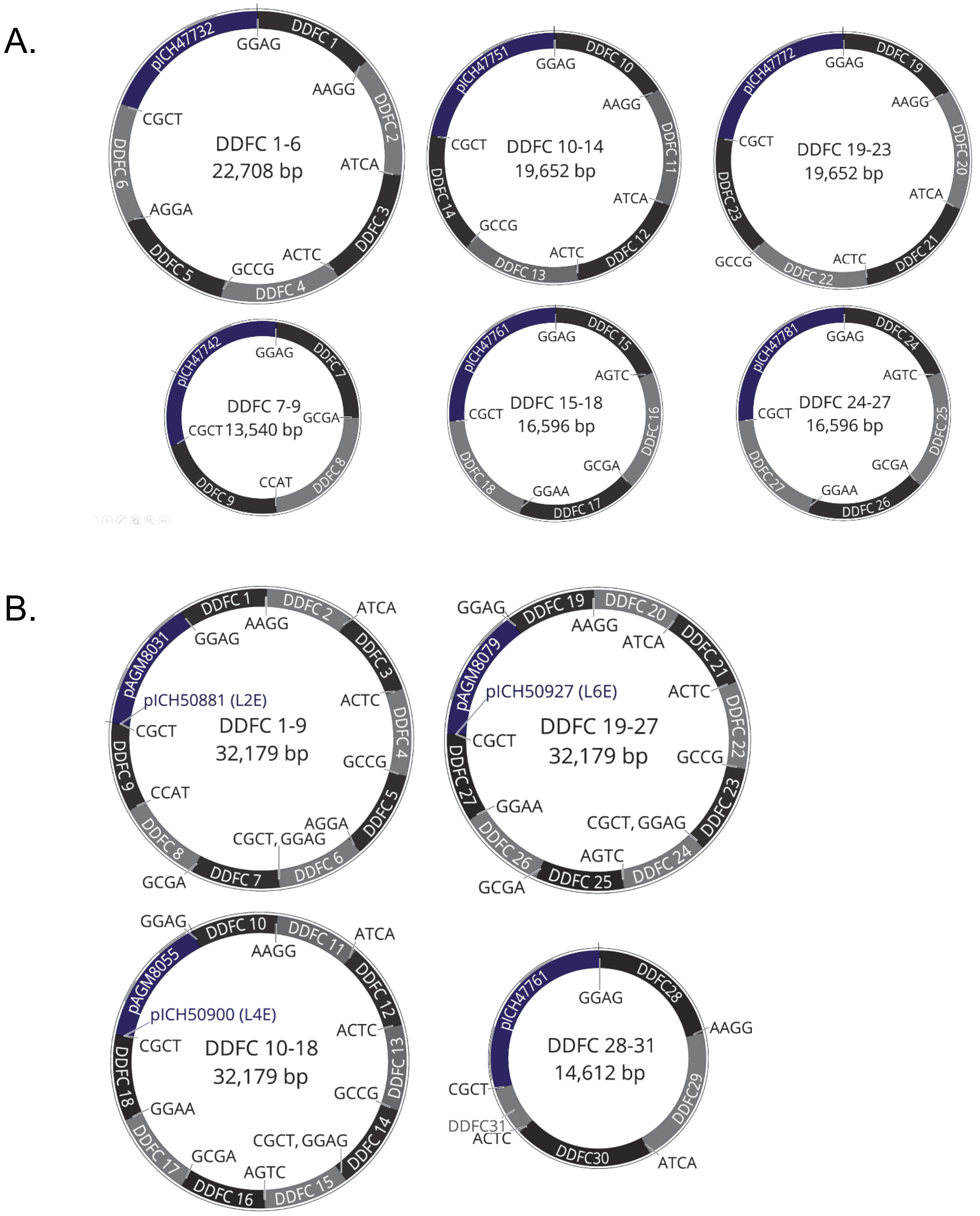


**Fig. S2. Maps of the DDFC plasmids**
**A.** Maps of the intermediate plasmids used to assemble from 3 to 6 sections each.
**B.** Maps of the 4 plasmids containing the 31 sectors of the DDFC file.

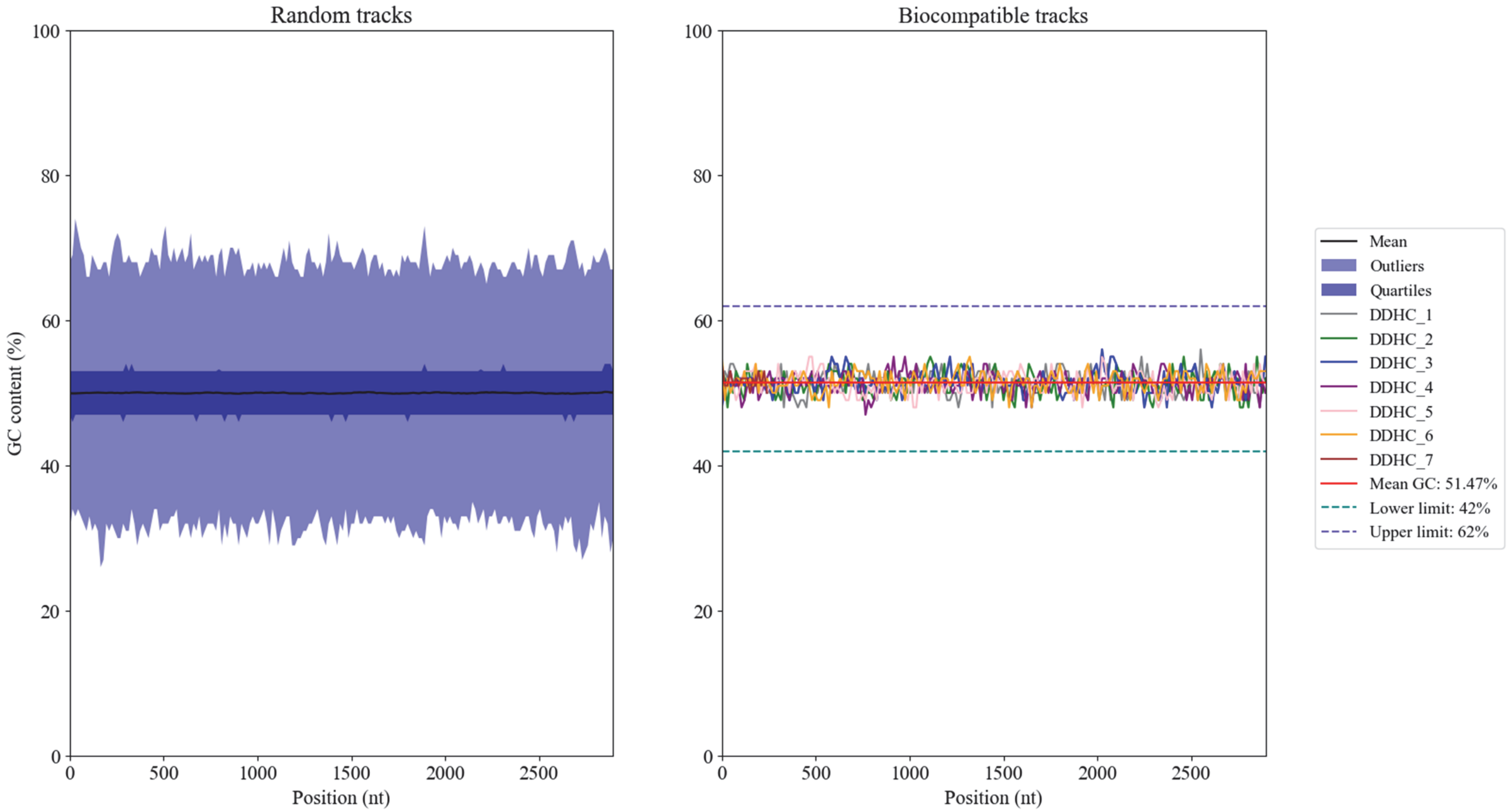


**Fig. S3. Distribution of GC content before and after RISE biocompatibilization**
A set of 1000 sequences was generated from the 7 binary chunks corresponding to the 7 sectors of DDHC by random draw using CGK conversion. The GC content was measured in windows/bins of 100 nucleotides and steps of 15 nucleotides for the 7000 sequences (left panel) or for the biocompatible sequence generated by RISE (right panel).

A.                                    B.

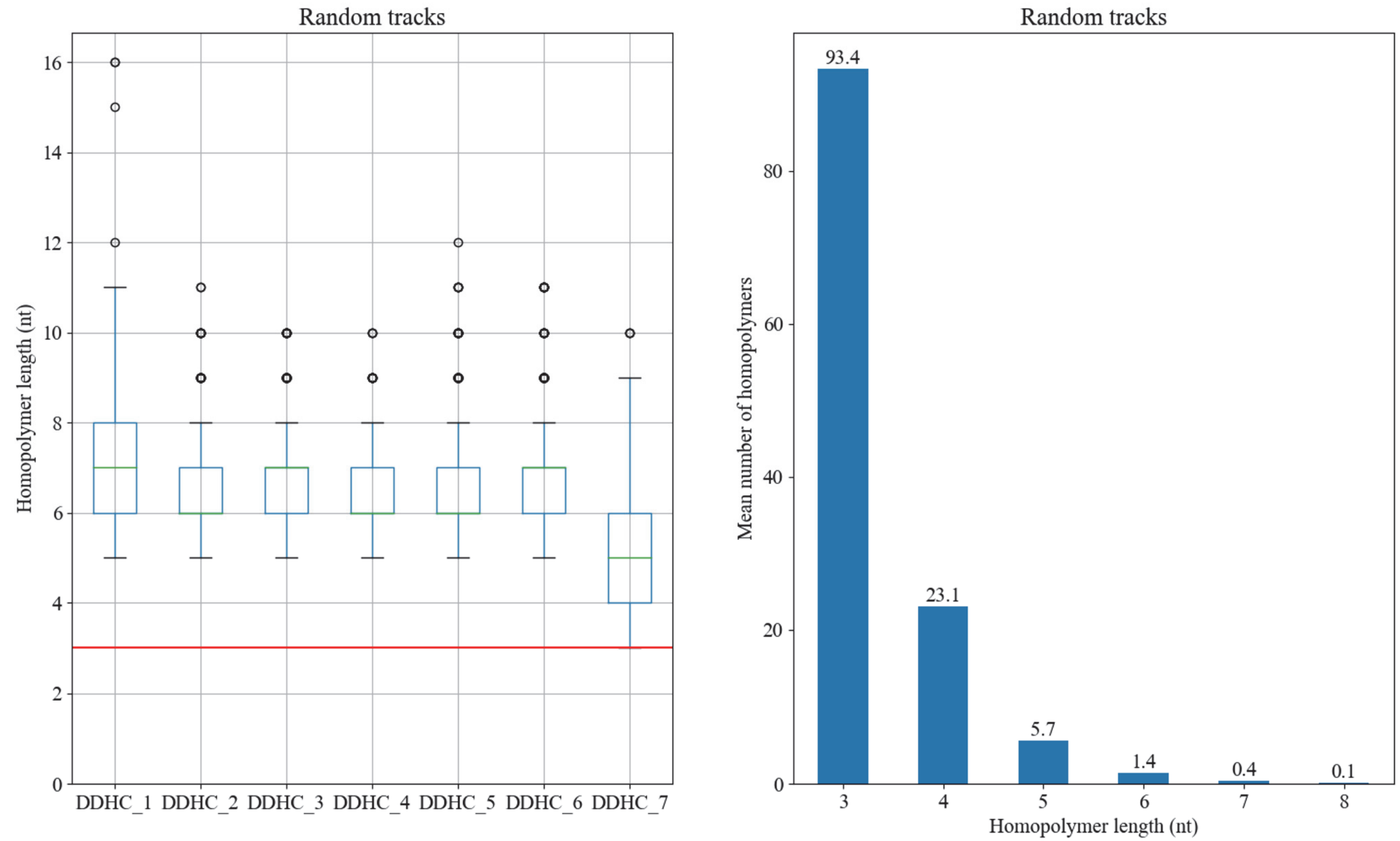


**Fig. S4. Distribution of homopolymer number and length before RISE biocompatibilization**
A set of 1000 sequences was generated from the 7 binary chunks corresponding to the 7 sectors of DDHC by random draw using CGK conversion. **A.** The maximum homopolymer length in each sequence is visualized with box and whisker plots showing the 10th (lower whisker), 25th (base of box), 75th (top of box) and 90th (top whisker) percentiles. Outliers are plotted as individual data points. The red line represents the maximum homopolymer size after RISE biocompatibilization.
**B.** The mean number of homopolymers in each of the 7000 sequences prior to biocompatibilization is plotted for different homopolymer lengths.

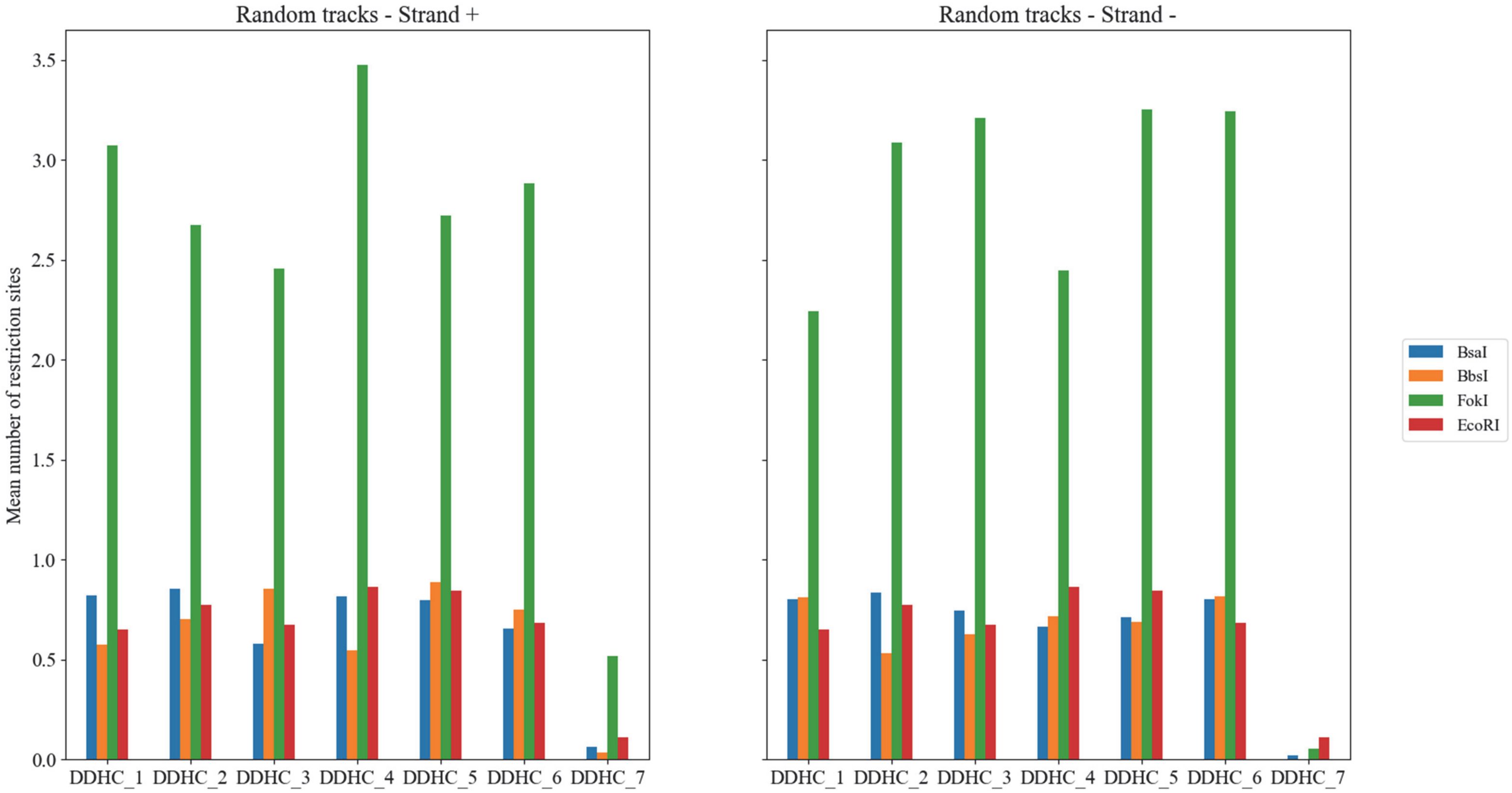


**Fig. S5. Distribution of specific restriction sites before complete removal by RISE**
A set of 1000 sequences was generated from the 7 binary chunks corresponding to the 7 sectors of DDHC by random draw using CGK conversion. The mean number of restriction sites in each of the 7000 sequences is plotted for BsaI, BbsI, EcoRI and FokI for the forward strand (left panel) and the reverse strand (right panel).

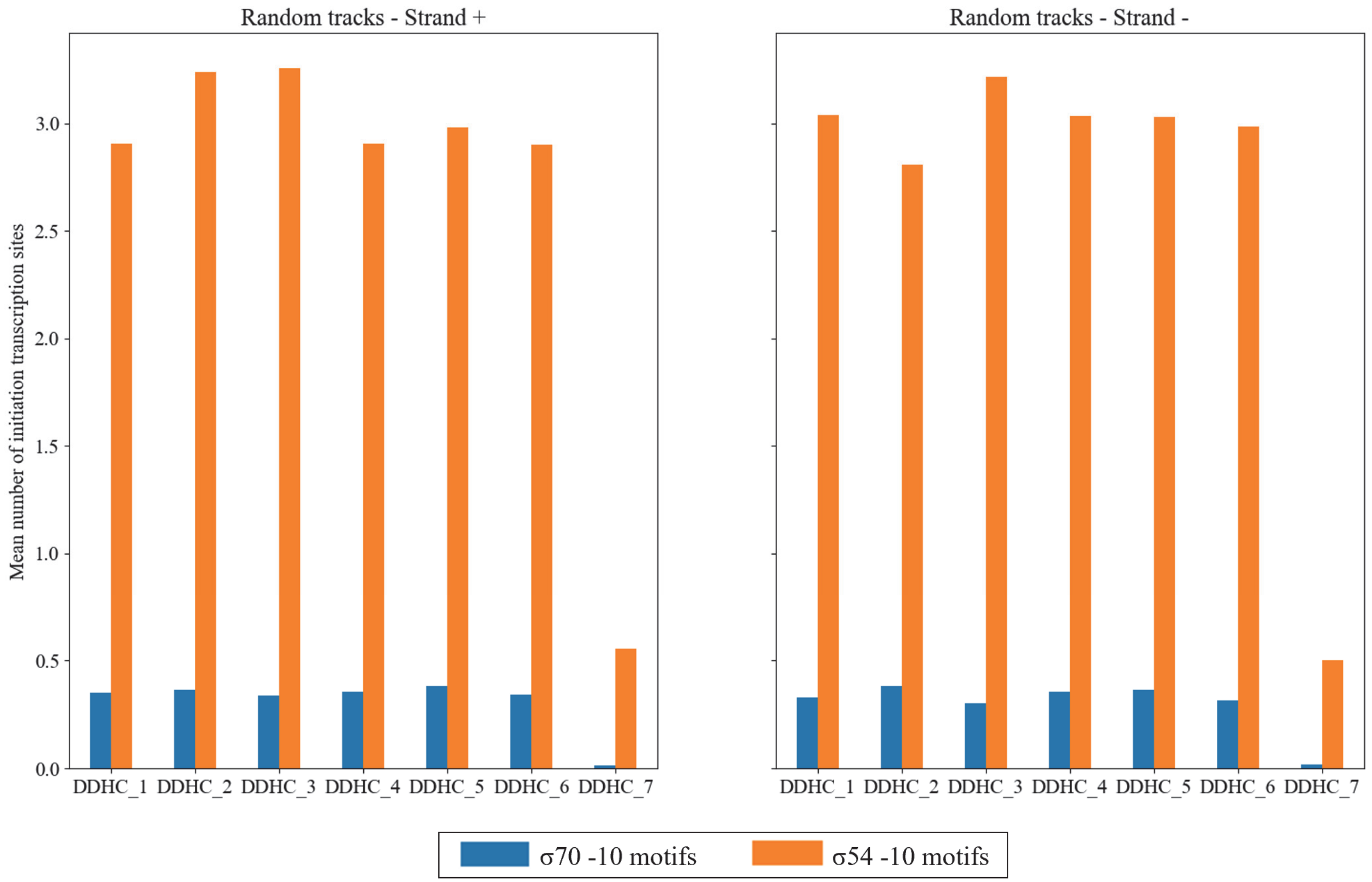


**Fig. S6. Transcription initiation site content before RISE biocompatibilization**
A set of 1000 sequences was generated from the 7 binary chunks corresponding to the 7 sectors of DDHC by random draw using CGK conversion. The mean number of $\sigma^{70}$ and $\sigma^{54}$ -10 motifs in each of the 7000 sequences is plotted for the forward strand (left panel) and the reverse strand (right panel).

A.

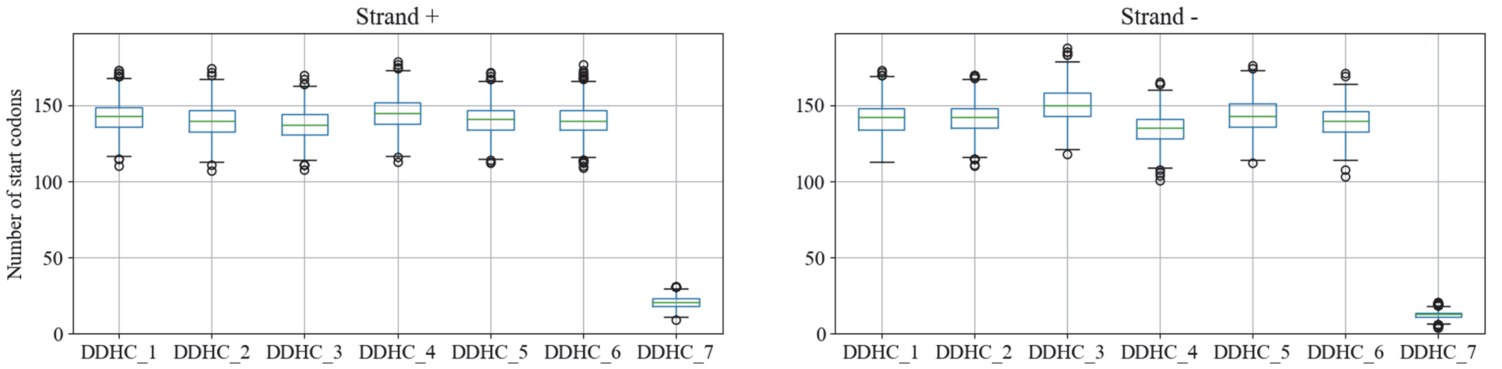


B.

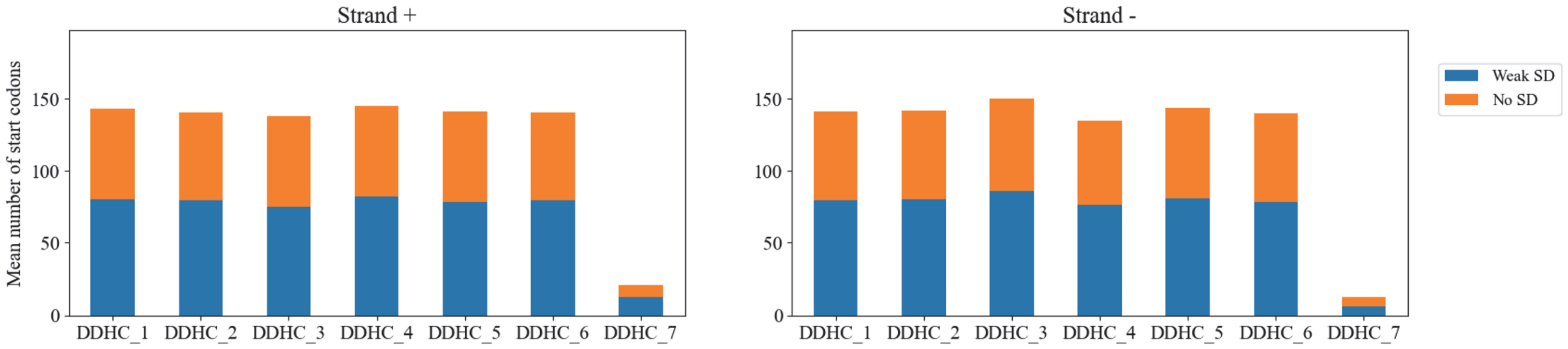


C.

| | DDHC_1 | DDHC_2 | DDHC_3 | DDHC_4 | DDHC_5 | DDHC_6 | DDHC_7 |
|---|---|---|---|---|---|---|---|
| Strong SD | 0.379 | 0.31 | 0.274 | 0.496 | 0.328 | 0.321 | 0.053 |
| Weak SD | 80.054 | 79.09 | 75.057 | 81.79 | 78.274 | 79.7 | 12.604 |
| No SD | 62.293 | 60.763 | 62.272 | 62.865 | 62.318 | 60.607 | 8.138 |

| | DDHC_1 | DDHC_2 | DDHC_3 | DDHC_4 | DDHC_5 | DDHC_6 | DDHC_7 |
|---|---|---|---|---|---|---|---|
| Strong SD | 0.33 | 0.421 | 0.371 | 0.332 | 0.417 | 0.389 | 0.02 |
| Weak SD | 79.675 | 79.941 | 85.508 | 75.875 | 80.821 | 77.766 | 5.865 |
| No SD | 61.407 | 61.399 | 64.367 | 58.238 | 62.461 | 61.64 | 6.664 |

**Fig. S7. Translation initiation sites before RISE biocompatibilization**
A set of 1000 sequences was generated from the 7 binary chunks corresponding to the 7 sectors of DDHC by random draw using CGK conversion. **A.** The total number of start codons (ATG, GTG, CTG) in each of the 7000 sequences is represented for the forward strand (left panel) and the reverse strand (right panel) as box and whisker plots showing the 10th (lower whisker), 25th (base of box), 75th (top of box) and 90th (top whisker) percentiles. Outliers are plotted as individual data points. **B.** The mean number of start codon with (blue) or without (orange) a weak Shine Dalgarno (SD) sequence for the forward strand (left panel) and the reverse strand (right panel).
**C.** Mean number of start codons with or without Shine Dalgarno sequences in each of the 1000 sequences of the 7 sectors.

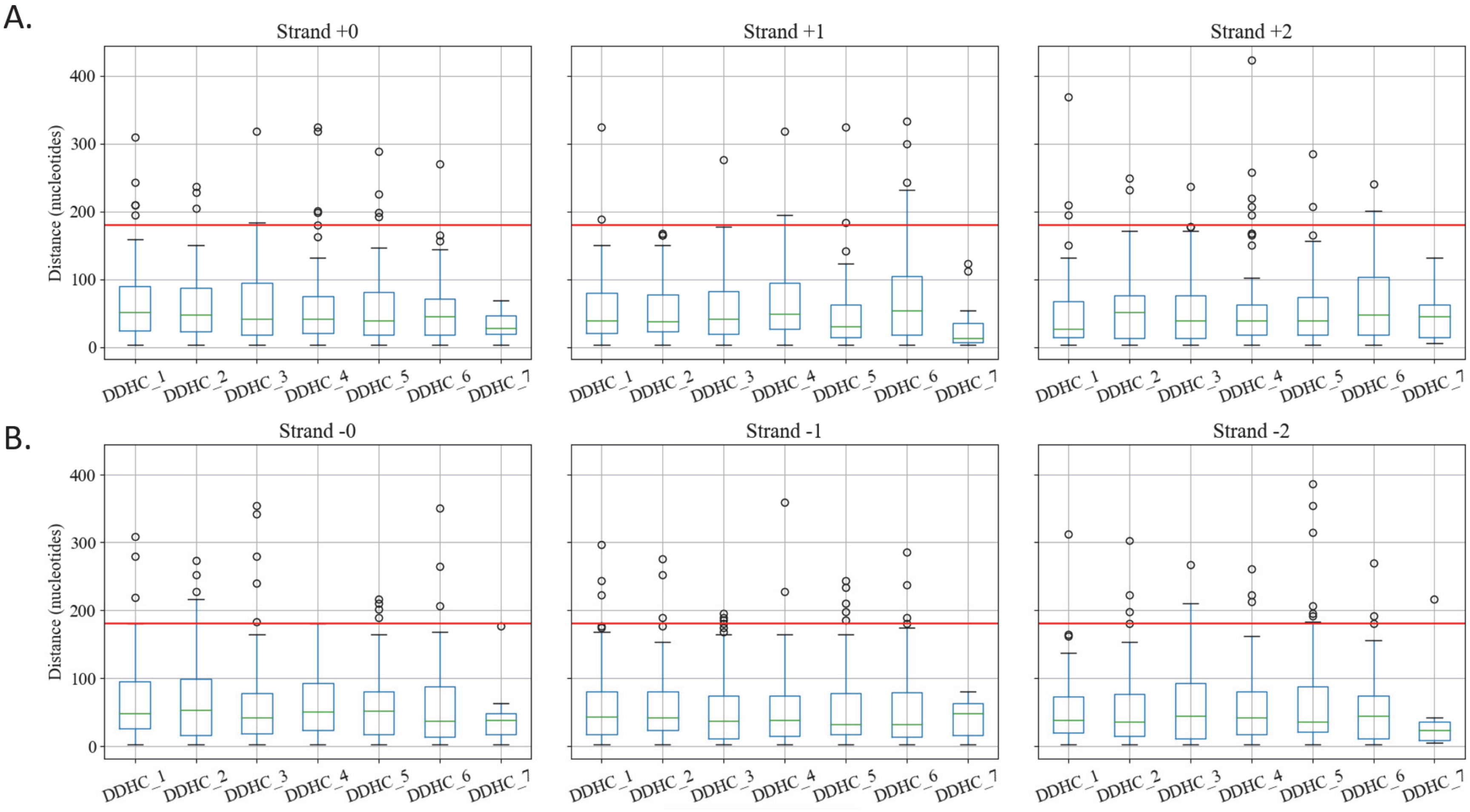


**Fig. S8. Distance between stop codons before RISE biocompatibilization**
A set of 1000 sequences was generated from the 7 binary chunks corresponding to the 7 sectors of DDHC by random draw using CGK conversion. The maximum interstop distance in the 3 frames of the forward strand (**A**) and the reverse strand (**B**) in each of the 1000 sequences of the 7 sectors is represented as box and whisker plots showing the 10th (lower whisker), 25th (base of box), 75th (top of box) and 90th (top whisker) percentiles. Outliers are plotted as individual data points. The red line represents the maximum interstop distance after RISE biocompatibilization.

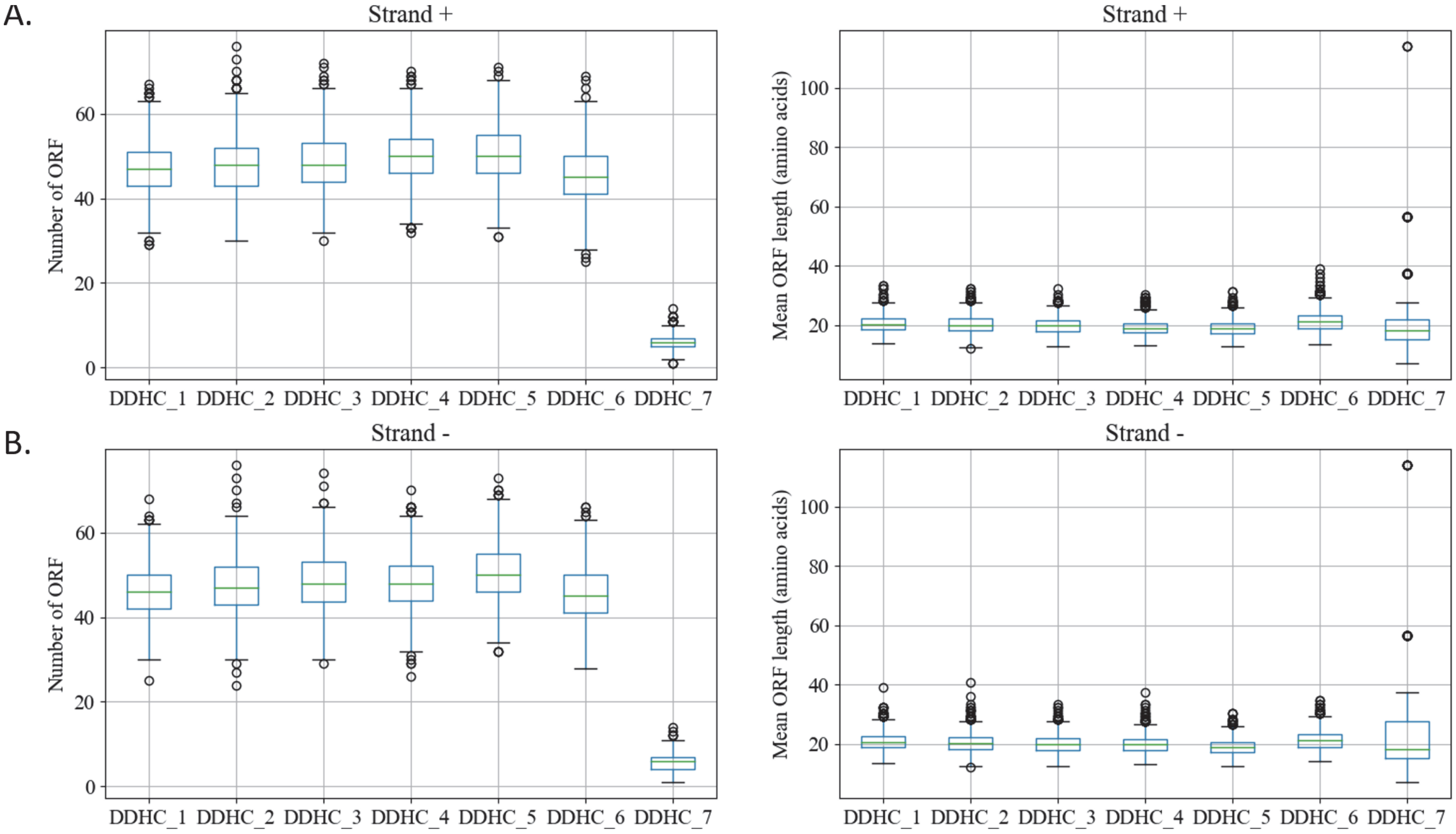


**Fig. S9. Number and frequency of open reading frames before RISE biocompatibilization**
A set of 1000 sequences was generated from the 7 binary chunks corresponding to the 7 sectors of DDHC by random draw using CGK conversion. Mean number (left panel) and mean length (right panel) of open reading frames in each of the 1000 sequences of the 7 sectors for the forward strand (**A**) and the reverse strand (**B**) represented as box and whisker plots showing the 10th (lower whisker), 25th (base of box), 75th (top of box) and 90th (top whisker) percentiles. Outliers are plotted as individual data points.

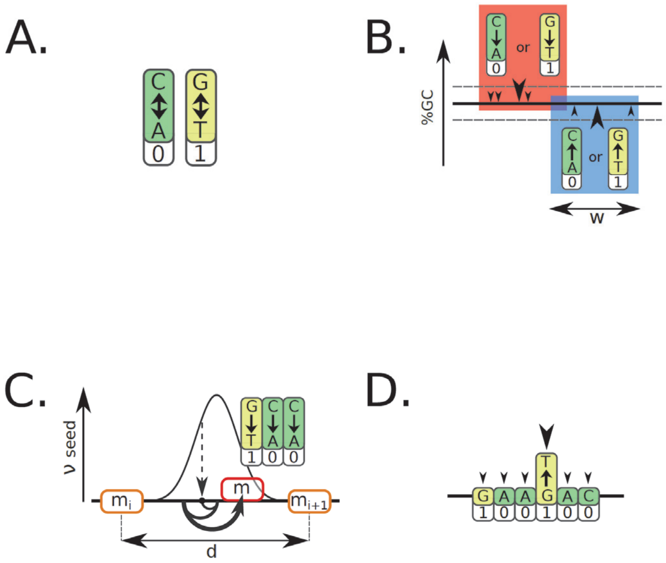


**Fig. S10. RISE processing for biocompatibilization**
**A.** During RISE, nucleotides can be swapped into their binary synonym without altering digital information. **B.** Sectors are sequentially adjusted for GC content. In a scanning window w, one random nucleotide is swapped to its higher or lower GC content synonym to reduce or increase the local GC content respectively. **C.** DNA motifs can be added by synonym swapping. Motif (m) density 1/d is increased by scanning and swapping for in-frame synonyms (circular arrow) *e.g.* stop codon TAA. The initial scanning position is selected between normally distributed positions with $\mu$=d/2 (dashed arrow). Addition is performed on the six translating frames. **D.** DNA motifs can be removed on both frames by randomly swapping one nucleotide of a forbidden motif *e.g.* BbsI restriction site GAAGAC.

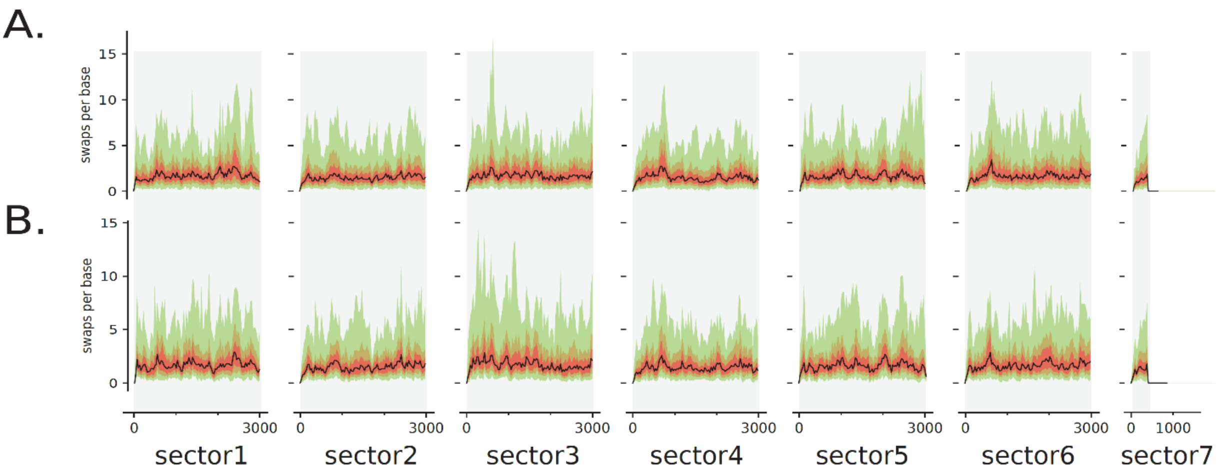


**Fig. S11. Distribution of the number of swaps over each sector during RISE biocompatibilization of DDHC**

**A.** Number of swaps per base during RISE starting with independent initial CGK random draws (N=100). **B.** Swaps during multiple RISE on the same initial random draw. Constraints are described in Fig. 2. Black line is the median of swaps/bin, red area is the first and third quartile of swaps distribution/bin, brown the 10th and 90th quantile and light green the minimum and maximum values.

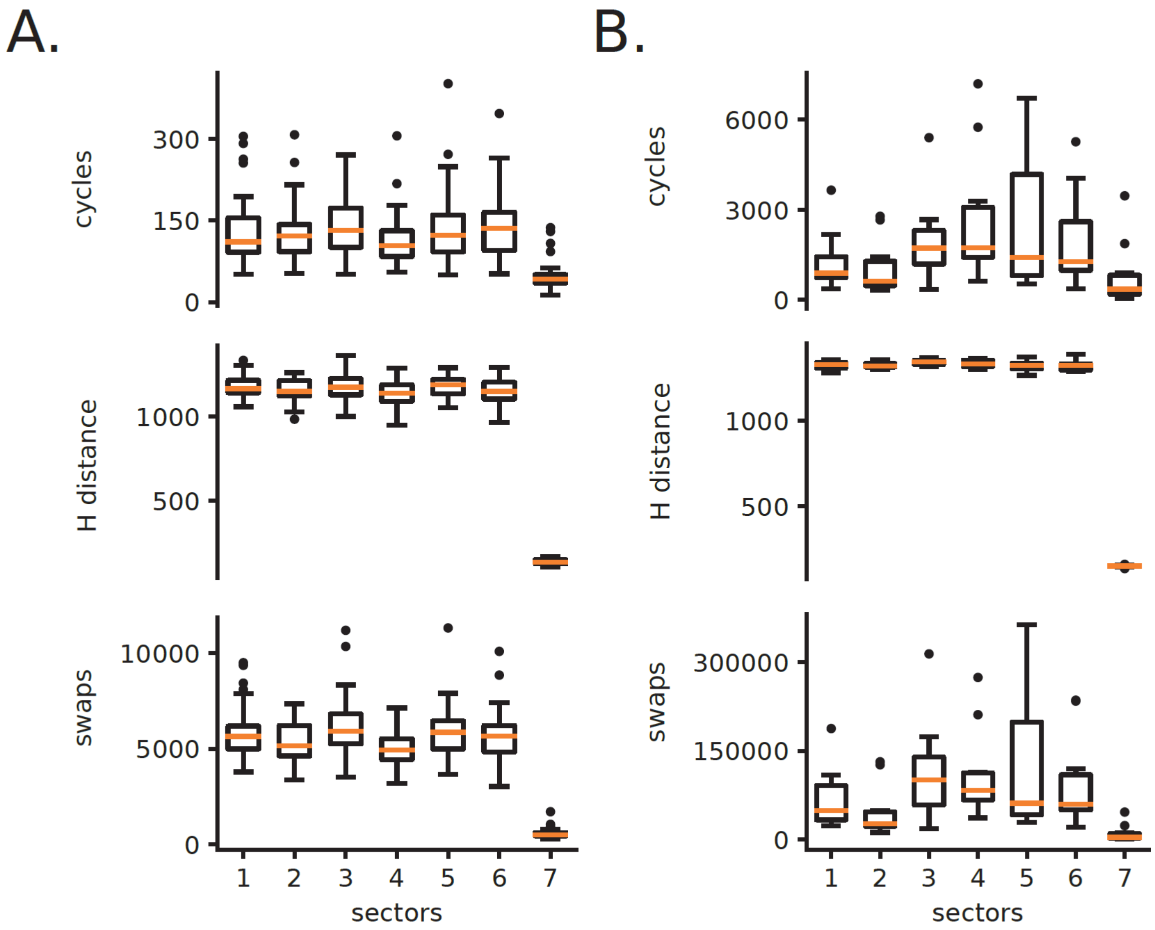


**Fig. S12. RISE robustness and efficiency**

**A.** Independent RISE simulations for one initial CGK random draw on the 7 sectors of DDHC (N=40).

Multiple RISE simulations on the same initial draw results in various final compatible sequences indicating that random driven processes of RISE allow various paths of compatibilization. For all runs and both files, approximately 1/3 of the initial nucleotides have been swapped to their CGK counterpart. The constraints are described in Fig. 2.

**B.** Detailed RISE simulations for 40 % GC (N=20).

The cycles indicate the number of RISE iterations. The Hamming distance (H distance) is computed between the initial and the final fully compatible sequence. Swaps are the number of nucleotide exchanges during all iterations. The box and whisker plots show the 10th (lower whisker), 25th (base of box), 75th (top of box) and 90th (top whisker) percentiles. Outliers are plotted as individual data points. All RISE simulations produced fully compatible sequences.

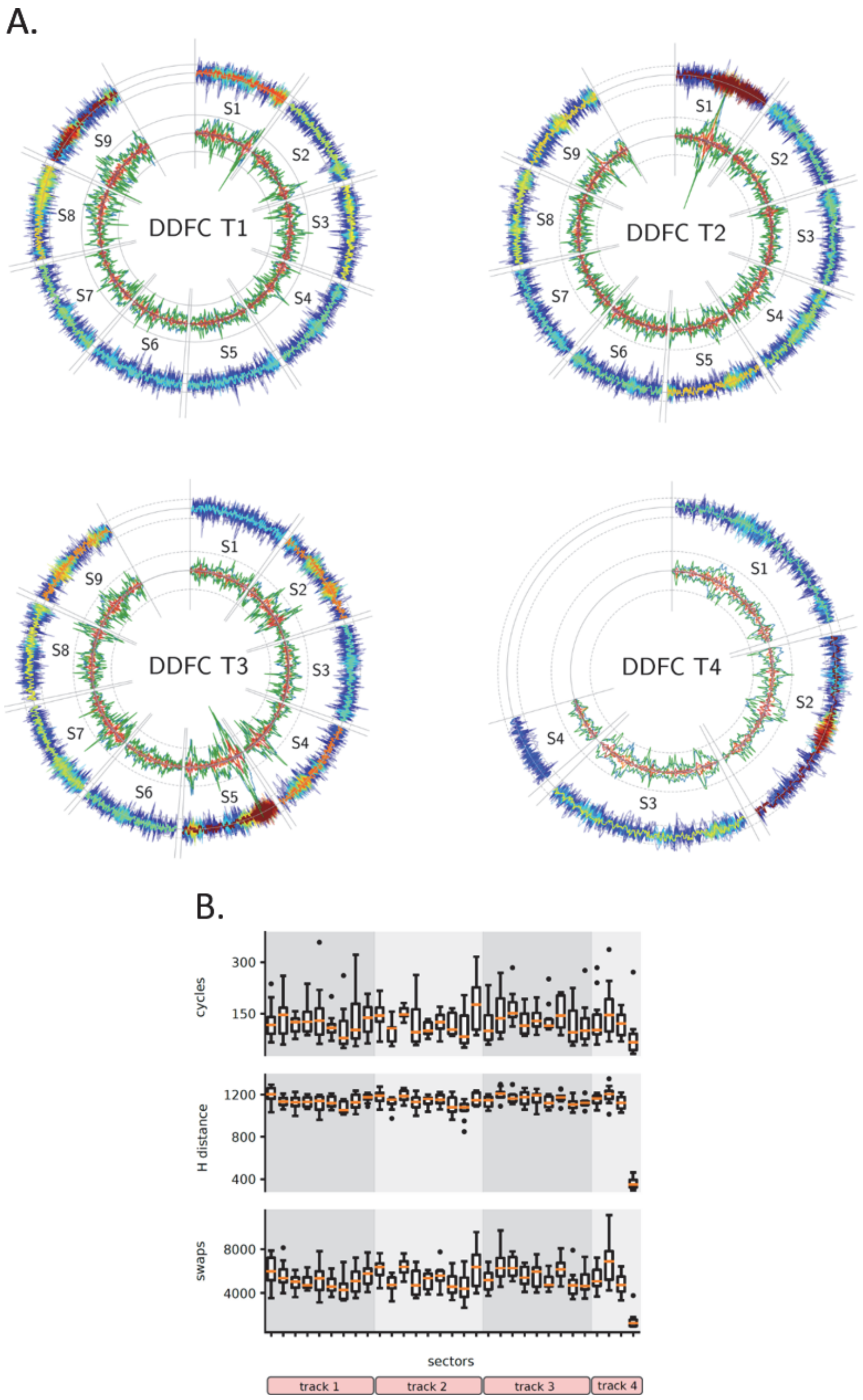


**Fig. S13. Biosafe and biocompatible encoding of DDFC**

Full biocompatibilization of the 31 sectors of DDFC by adding or removing motifs on both strands and adjusting GC content. **A.** Number and nature of swap positions during RISE (inner circle, mean swaps in w = 30 bp) and GC content evolution (outer circle). Colors as in Fig. 3.

**B.** RISE efficiency on the 31 sectors of DDFC (D) (N=40). The box and whisker plots show the 10th (lower whisker), 25th (base of box), 75th (top of box) and 90th (top whisker) percentiles. Outliers are plotted as individual data points. Hamming (H) distance is computed between the initial and final fully compatible sequence.

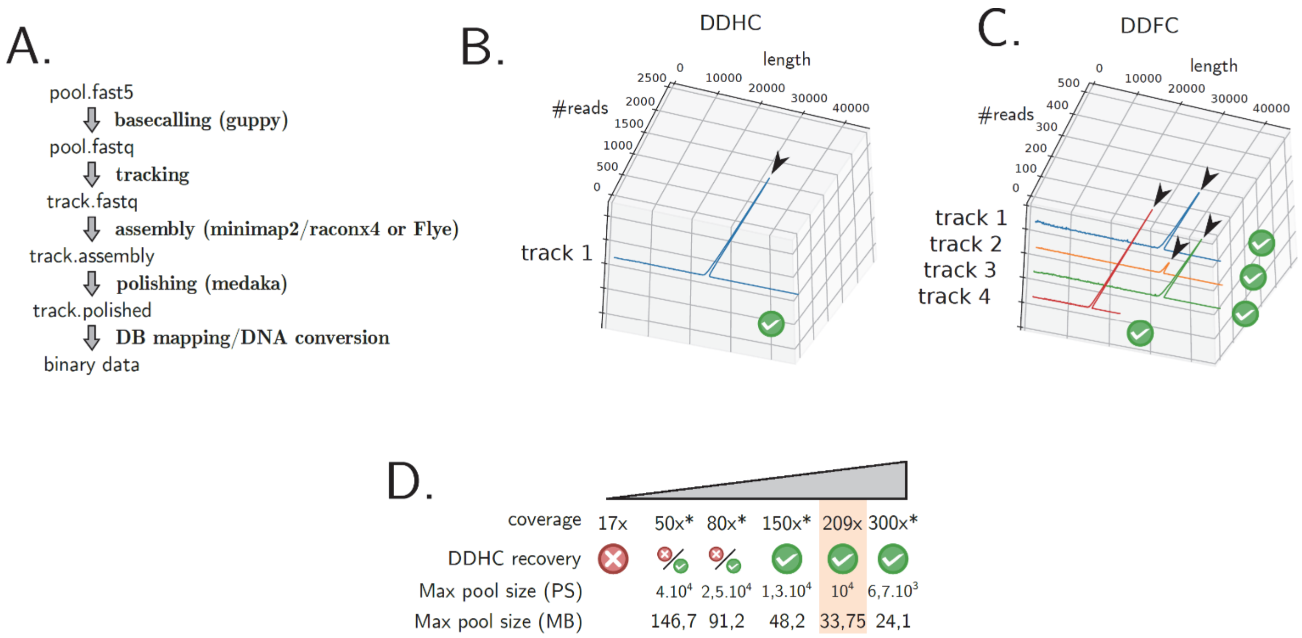


**Fig. S14. Nanopore sequencing of the DDHC and DDFC DNA drives**
**A.** Overall process of DNA Drive sequence recovery after Nanopore sequencing.
**B.** and **C.** Read size distribution after DNA Drive sequencing (DDHC and DDFC DNA Drive respectively). Each line corresponds to one track. Black arrow indicates expected track length. Green check mark indicates complete data retrieval.
**D.** Required coverage for complete data retrieval using Nanopore MinIon.
Experimental and simulated sequence coverage for DDHC is indicated. For experimental coverage, DDHC Track 1 is diluted into DDFC Track 4 prior to sequencing. Simulated coverage (*) is obtained by subsampling random experimental DDHC reads. Reads are processed as described in panel A using Flye and minimap2/racon for draft assembly. Green check mark and red cross indicate complete or incomplete data retrieval of DDHC, respectively. For intermediate coverage (50x and 80x), complete data retrieval was possible with Flye software but not with minimap2/racon. Max pool size (PS) indicates the maximum number of tracks per pool for a 50 Gb Minion run. The corresponding max pool size (in MB) for B3000S1I25TS9 tracks is indicated.
